# Ionizing radiation induces stem cell-like properties in a caspase-dependent manner in *Drosophila*

**DOI:** 10.1101/316497

**Authors:** Shilpi Verghese, Tin Tin Su

## Abstract

Cancer treatments including ionizing radiation (IR) can induce cancer stem cell-like properties in non-stem cancer cells, an outcome that can interfere with therapeutic success. Yet, we understand little about what consequences of IR induces stem cell like properties and why some cancer cells show this response but not others. In previous studies, we identified a pool of epithelial cells in *Drosophila* larval wing discs that display IR-induced stem cell-like properties. These cells are resistant to killing by IR and, after radiation damage, change fate and translocate to regenerate parts of the disc that suffered more cell death. Here, we addressed how IR exposure results in the induction of stem cell-like behavior, and found a requirement for caspase activity. Unexpectedly, this requirement was mapped to the regenerative cells, suggesting a non-apoptotic role for caspases in the induction of stem cell-like behavior. We also performed a systematic probing of different regions of the wing disc by lineage tracing, in order to identify additional pools of cells with IR-induced regenerative properties. We identified two new populations of such cells. Unlike the original pool that helps regenerate the disc, the new pools of cells undergo abnormal regeneration to produce an ectopic, supernumerary wing disc. We also identified cells that lack the ability to display IR-induced regenerative behavior. Identification of different cell populations with different IR-induced regenerative potential will allow us to probe the molecular basis for these differences in the future.

**AUTHOR SUMMARY:** Ionizing Radiation (IR), alone or in combination with other therapies, is used to treat an estimated half of all cancer patients. Yet, we understand little about why some tumors cells respond to treatment while others grow back (regenerate). We identified specific pools of cells within a *Drosophila* organ that are capable of regeneration after damage by IR. We also identified what it is about IR damage that allows these cells to regenerate. These results help us understand how cells regenerate after IR damage and will aid in designing better therapies that involve radiation.

## INTRODUCTION

Regeneration is essential to tissue homeostasis and health. Conversely, regeneration of tumors after treatment leads to tumor recurrence and treatment failure. Understanding mechanisms that underlie regeneration is therefore important not only for understanding basic biology but also for optimizing treatment of diseases like cancer. Our understanding of regeneration has benefited immensely from experimental systems with dedicated stem cells that form the cellular basis for regeneration. Examples include regeneration of vertebrate gut and *Drosophila* intestine [1–3]. Tissues also regenerate despite the lack of a dedicated stem cell pool. A prime example is the vertebrate liver, which regenerates by proliferation of the surviving cells of each cell type [4–6]. If proliferation of hepatocytes is blocked during liver regeneration, however, biliary epithelial cells can dedifferentiate, proliferate and re-differentiate into hepatocytes [4–6]. Such plasticity has been documented in other mammalian organs [7–9], and in some models of amphibian limb and fish fin regeneration [10]. This report addresses the molecular basis for cell fate plasticity during regeneration using *Drosophila* larval cells as a model.

*Drosophila* larval imaginal discs are precursors of adult organs. Imaginal discs lack a dedicated stem cell pool yet can regenerate fully even after surgical ablation of 25% of the disc, after genetic ablation of a disc compartment (e.g. the anterior compartment), or after exposure to doses of ionizing radiation (IR) that kills about half of the cells [11, 12]. We recently identified a previously unknown mode of regeneration in *Drosophila* larval wing discs, whereby epithelial cells acquire stem cell-like properties during regeneration after damage by IR [13]. These properties include resistance to killing by IR, the ability to change cell fate, and the ability to translocate to areas of the wing disc with greater need for cell replenishment. The ability to behave like stem cells in response to IR is limited to certain cells within the continuous epithelium of the wing disc. Specifically, a subset of future hinge cells is protected from IR-induced apoptosis by the action of STAT92E (*Drosophila* STAT3/5, to be called ‘STAT’ hereafter) and Wg (Dm Wnt1)-mediated repression of pro-apoptotic gene *reaper*. These hinge cells change their fate and translocate to the pouch region that suffers more apoptosis and participate in regenerating the pouch. Without IR, these cells differentiate into the adult wing hinge, indicating the cell fate plasticity is IR-induced.

In above-described studies, regeneration of the pouch by the hinge was observed in nearly all irradiated discs [13, 14]. In about 20% of irradiated discs, we observed, in addition, abnormal regeneration that produced an ectopic wing disc [14]. Ectopic discs were wing discs based on staining for the protein markers Ubx and Wg, and are composed of an ectopic pouch and an ectopic hinge [14]. Ectopic discs were neither duplications (e.g. not pouch-to-pouch) nor transdeterminations (e.g. not leg-to-wing) described in classical studies of regeneration after surgical ablation [15]. Our efforts to dissect the cellular origin for the ectopic discs showed that cells of the hinge that regenerate the pouch are unlikely to be responsible for ectopic discs [14]. Therefore, we hypothesized that there are additional pools of cell in the wing disc that show stem cell like properties after IR damage by participating in abnormal regeneration to produce an ectopic wing disc.

Here, we report the mapping of cell lineages during regeneration of *Drosophila* larval wing discs following damage by X-rays, a type of IR. To express lineage tracers, we used FlyLight GAL4 drivers that display simple expression patterns because they use small (^~^3kb) enhancers from various genes [16]. In addition to the subset of future hinge cells we previously identified as capable of behaving like stem cells [13], two more cell populations, in the notum and in the dorsal-posterior hinge/pleura region, were found to show this potential. While the previously identified hinge cells are responsible for normal regeneration to restore the wing disc, the newly identified regenerative cells undergo abnormal regeneration to produce ectopic discs. Cells of the pouch, we find, lack the capacity to acquire stem cell like properties and do not change fate or translocate. Of possible consequences of X-ray exposure, we identified caspase activity, as necessary for the hinge cells to acquire regenerative properties, and further localize this requirement to the regenerative cells.

IR-induced stem-ness in *Drosophila* parallels the increasingly appreciated ability of cancer treatments including IR to induce stem cell-like properties in non-stem cancer cells [17–20]. Furthermore, within the continuous single layer columnar epithelium of the wing disc, only some cells respond to IR by exhibiting stem cell-like properties. This parallels how only a small subset of irradiated cancer cells exhibit stem cell-like properties. IR, alone or in combination with other therapies, is used to treat an estimated half of all cancer patients. Yet, we understand little about what consequences of IR induces stem cell like properties and why some cancer cells show this response but not others. This report describes similar phenomena in *Drosophila*, making it possible to identify and study the responsible genes in the future.

## RESULTS

We are using the published G-trace system to monitor cell lineages in the larval wing discs [21]. In this system, GAL4 drives the expression of UAS-RFP (real time marker) and UAS-FLP recombinase, which causes a recombination event to produce stable GFP expression (lineage marker). Thus, even if cells change fate and lose RFP expression, their clonal descendants would be marked with GFP (GFP^+^ RFP^-^). We used G-trace to test a collection of GAL4 drivers, each active within a different subset of cells in *Drosophila* 3^rd^ instar larval wing discs.

### Optimization of lineage tracing protocol

30A-GAL4 was used in our recent studies and is active in a subset of hinge cells ([13, 14]; reproduced in Fig 1A-C, J, L) and a small number (<10) of notum cells (arrows in Fig 1 J, L). The dynamics of 30A-GAL4 activity is such that cells expressing it show stable lineage; very few cells were GFP RFP. In contrast, another hinge driver, FlyLight R73G07-GAL4 produced GFP^+^ cell in most of the disc including the pouch, the hinge, and most of the notum, even though RFP is restricted to the hinge in 3^rd^ instar wing discs (Fig 1D-F). R73G07 is a 3028 bp enhancer from the *zfh2* locus and is apparently active in most cells of the wing disc before becoming restricted to the hinge. Zfh2 is a transcription factor important for wing development [22]. Temporal restriction of R73G07-GAL4 activity with its repressor GAL80^ts^ according to the temperature shift protocol shown in Fig 1M confined GFP to the hinge and the pouch (Fig 1I). Increasing larval age from 4-5 days after egg laying (AED, Fig 1M) to 5-6 days AED before temperature shift to induce GAL4 did not reduce the GFP^+^ domain any further. Further aging the larvae beyond 5-6 days AED before inducing GAL4 may help eliminate GFP in the pouch and restrict it to the hinge, but this schedule is incompatible with our goal because we need to monitor regeneration for 72 h after IR before losing the larvae to pupariation.

**Figure 1.**
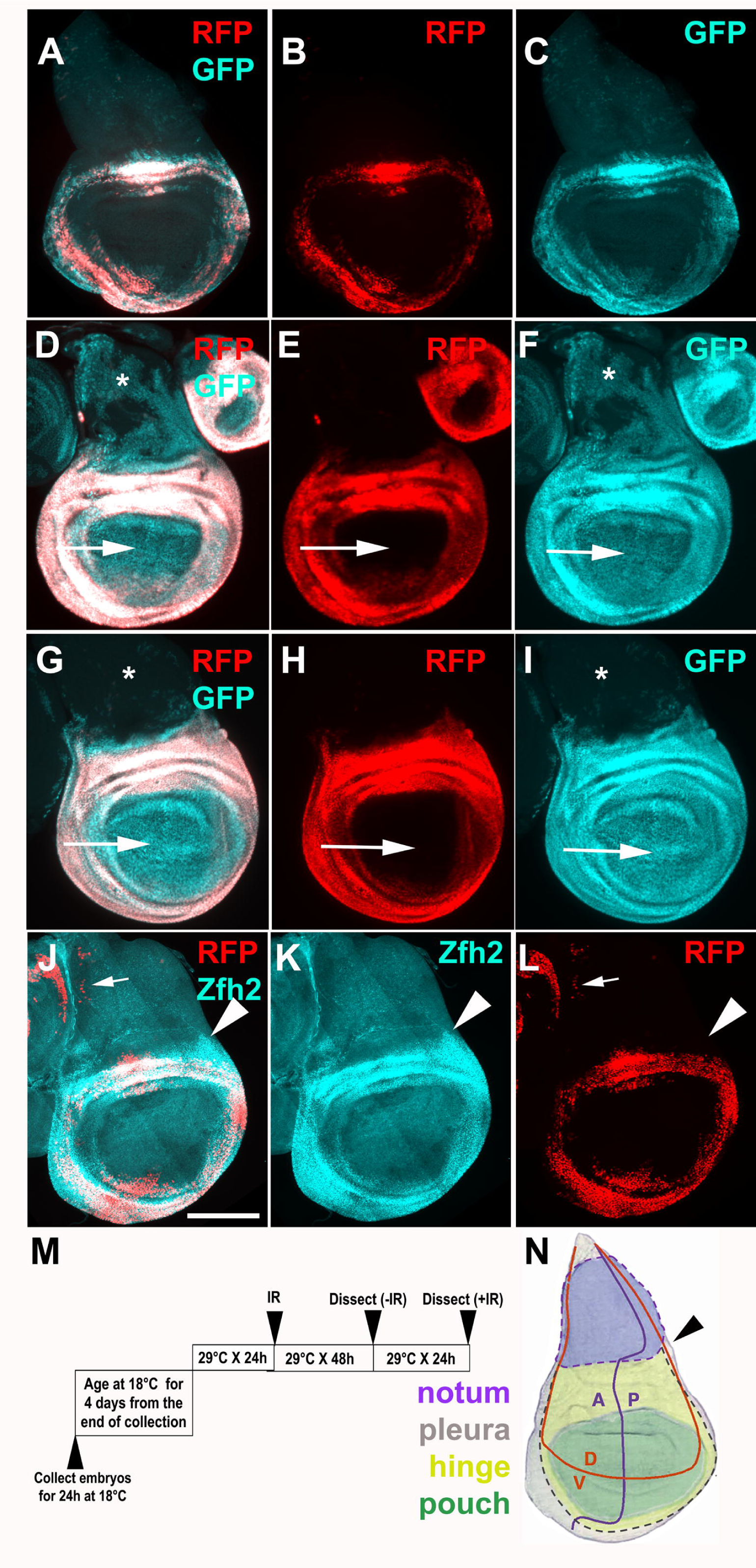
Lineage tracing with two GAL4 drivers that are active in the hinge. Wing discs were removed from 3^rd^ instar larvae without irradiation, fixed and imaged for RFP/GFP. The discs in J-K were also stained with an antibody to Zfh2. All discs are shown with anterior (A) left and dorsal (D) up as in (N). Scale bar = 100 microns. (A-C) 30A-GAL4 drives UAS-RFP (real time marker) and GFP (lineage marker) (D-I) R73G07-GAL4 drives UAS-RFP (real time marker) and GFP (lineage marker). Restricting GAL4 activity with GAL80^ts^ eliminated the expression of GFP in the notum (^*^) in (G-I). (J-L) Zfh2 antibody staining and 30A>RFP expression. Zfh2-expressing cells outside the 30A domain are indicated with arrowheads. (M) The temperature shift protocol. Embryos were collected at 18°C for 24 h and cultured at 18°C until 4-5 days after egg laying, reaching late 2^nd^ instar. Larvae were shifted to 29°C for 24 h to reach early 3^rd^ instar before irradiation with 0 or 4000 R of X-rays. Discs were dissected 48 h later (for –IR controls) or 72 h after IR (+IR samples) because IR delays development. (N) The fate map with dotted lines added to indicate the pleura. Arrowhead points to the region that is Zfh2-expressing but outside the 30A domain in (J-L). Figure modified from [44]. The genotypes were: (A-C) 30A-GAL4, UAS-G-trace (see S1 Table for G-trace genotype)/SM5 (D-F) UAS-G-trace/+; R73G07-GAL4/+ (G-I) UAS-G-trace/+; R73G07-GAL4/tub-GAL80^ts^ (J-L) 30A-GAL4, UAS-G-trace /+; tub-GAL80^ts^ /+

These results illustrate that while some GAL4 drivers show stable lineage expression and could be used to monitor fate changes after irradiation, others show lineage changes without IR. This was confirmed using fifteen additional FlyLight GAL4 drivers (S1 Fig and figure legend). Therefore, we used GAL80^ts^ and the protocol shown in Fig 1M in all subsequent experiments with all GAL4 drivers, even if their lineages are stable as in the 30A-GAL4 example. We selected FlyLight drivers for their apparently exclusive expression in the disc region of interest. We find, however, that most show additional expression elsewhere in the disc, which would make interpretation of lineage results difficult. Therefore, in subsequent experiments, we used only the drivers that are expressed exclusively in cells of interest. Some GAL4 drivers are active in the cells of the peripodial membrane that covers the wing disc epithelium on the apical side, and in wing-disc associated tracheal cells on the basal side. In such cases, peripodial cells and tracheal cells can be identified based on their larger nuclear size compared to columnar epithelial cells and on their location in optical sections that book-end the columnar epithelium (S2 Fig). Our analyses focus on the columnar epithelium by excluding other optical sections.

Antibody staining shows that Zfh2 protein expression resembles R73G07-GAL4>RFP expression (compare Fig 1H and K). In contrast, 30A>RFP is expressed in only a subset of these cells (compare Fig 1K and L). Of relevance to subsequent sections is the expression of R73G07-GAL4 but not 30A-GAL4 in the dorsal-posterior region of the hinge and the pleura (arrow heads in Fig 1J-L, N).

### Cells of the dorsal-posterior hinge and pleura change fate and translocate into the notum

In our published studies of lineage tracing after irradiation, 30A-GAL4 expressing hinge cells translocated to the pouch but showed little movement dorsally towards the notum ([13, 14] and reproduced in Fig 7). We used 4000R of X-rays in this and all other experiments. This level of IR kills more than half of the cells but the discs could still regenerate to produce viable adults [11, 12]. In contrast to 30A-GAL4, R73G07-GAL4>G-trace experiments show GFP^+^ cell populations that extend from the hinge dorsally along both the anterior and posterior margins of the wing disc (Fig 2 I-P). The extending GFP^+^ cell population is contiguous with both the hinge (arrowheads) and the pleura (arrows in Fig 2F-H and Fig 2J-L; see fate map in Fig 1N). Some GFP^+^ cells in the notum lack RFP (for example, Fig 2H) while others express RFP (for example, Fig 2L). In discs that show an ectopic disc (Fig 2M-P), many cells of the ectopic disc are GFP^+^ RFP^+^. To better understand the source of GFP^+^ cells in the notum, we repeated the experiment but analyzed the discs at different times after IR.

**Figure 2.**
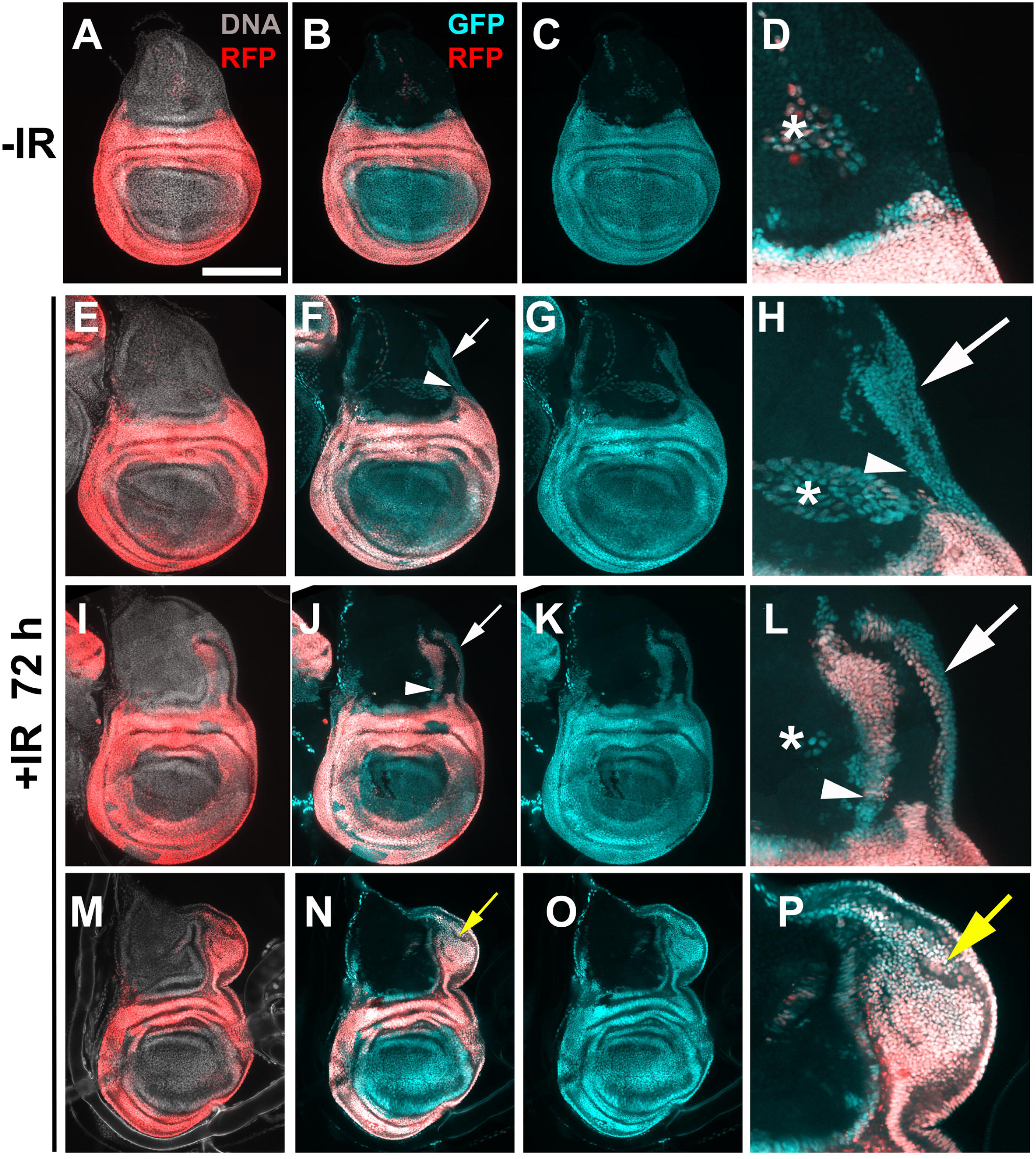
Posterior-dorsal hinge cells translocate towards the notum after IR. Larvae of the genotype UAS-G-trace/+; GAL80^ts^ / R73G07-GAL4 were treated as in Fig 1M. Wing discs are removed, fixed and imaged for RFP/GFP. All discs are shown with anterior (A) left and dorsal (D) up. Scale bar = 33 microns in D, H, L and P and 100 microns in the rest of the panels. Arrowheads = GFP^+^ cell populations in the notum that are contiguous with the hinge. Arrows = GFP^+^ cell populations in the notum that are contiguous with the pleura. ^*^ = cells outside the columnar epithelial layer that express G-trace.

We analyzed *a single cohort* of R73G07>G-trace larvae at 24, 48 and 72 h after IR, in two independent time course experiments (Fig 3). At 24 h after IR, GFP^+^ RFP^+^ cells are seen spreading from the hinge (arrowhead) and the pleura (arrow) into the notum (Fig 3A-B, magnified in C). Such translocation is not seen in -IR samples (compare Fig 3C to Fig 2D). The retention of RFP in these cells could be due to the persistence of GAL4, RFP, or both proteins. The half-life of GFP is 26 h in mammalian cells [23]. If the half-life of RFP in the wing disc is similar, these cells could have terminated RFP expression but still have RFP protein. The movement of hinge/pleura cells is seen along both the anterior and the posterior disc margins, but not in the central portion of the disc (Fig 3B). Such translocation of the hinge/pleura cells was seen in most discs (50/58 in two independent experiments). Of the remainder, three resembled –IR controls, that is, apparently without cell movements. The other five resembled what is described next for the 48 h time point.

**Figure 3.**
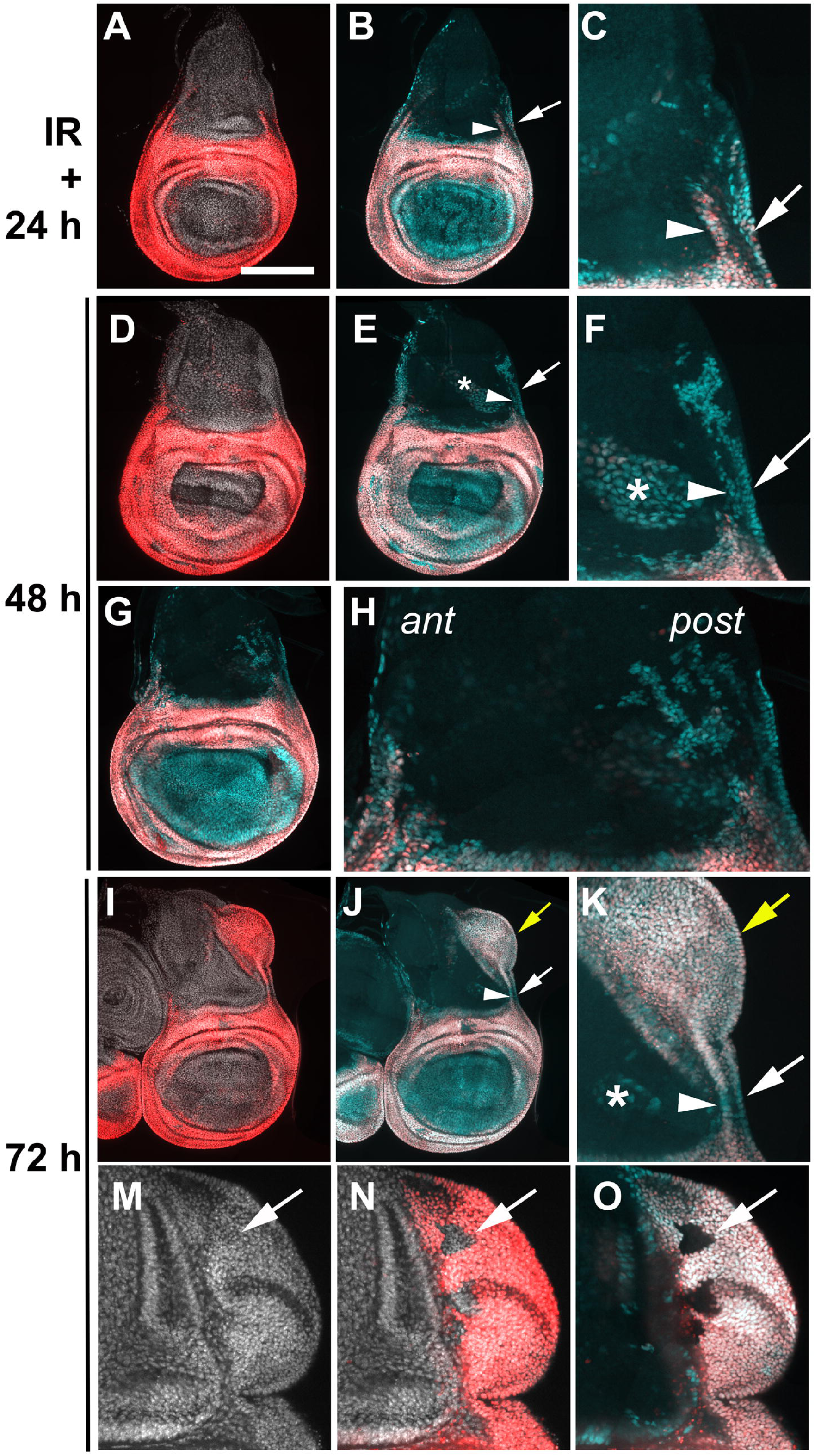
R73G07-GAL4>G-trace in a time course. Larvae of the genotype UAS-G-trace/+; GAL80^ts^ / R73G07-GAL4 were treated as in Fig 1M. Wing discs are removed, fixed and imaged for RFP/GFP at 24, 48 or 72 h after IR. The discs were also stained for DNA. All discs are shown with anterior (A) left and dorsal (D) up. Magnified panels show the relevant portion of the disc preceding it, except for M-N that shows the ectopic disc from a second 72 h disc. Scale bar = 33 microns in C, F, H, K and O, and 100 microns in the rest of the panels. Arrowheads = GFP^+^ cell populations in the notum that are contiguous with the hinge. Arrow = GFP^+^ cell populations in the notum that are contiguous with the pleura. ^*^ = cells outside the columnar epithelial layer that express G-trace.

At 48 h after IR, about one fourth of discs show movement of hinge and pleura cells into the notum, similar to the 24 h disc shown in Fig 3A-C (7/26, two independent experiments). In the majority of the discs (17/26, two independent experiments), the number of GFP^+^ cells in the notum increased, they are found deeper (more dorsal) in the notum, and most of these cells lacked RFP (Fig 3E-H). We interpret this to mean that cells continued to translocate from the R73G07-GAL4 domain and many of these have now lost their hinge fate as indicated by the lack of RFP. As in the 24 h discs, GFP^+^ cell population in the notum appear contiguous with both the hinge (arrowhead) and the pleura (arrow, Fig 3 E, magnified in F). Moreover, GFP^+^ cells in the notum were more numerous in the posterior half (post) than in the anterior half (ant) in some of the discs (Fig 3G, magnified in H), which may explain the finding that ectopic discs seen at 72 h after IR always appear along the posterior wing margin (e.g. Fig 2M-P; [14]). The remaining two discs show RFP^+^ GFP^+^ cells deep in the notum and far from the hinge, similar to what is shown in Fig 2J.

At 72 h after IR, we again see different classes as previously seen in Fig 2. The majority resemble the ones in Fig 3D-H, with GFP^+^ RFP^-^ cells in the notum (46/65, two independent experiments). About 15% (10/65, two independent experiments) of the discs show GFP^+^ RFP^+^ cells deep in the notum and far from the R73G07-GAL4 domain. In some of these discs, GFP^+^ RFP^+^ cells are contained within the notum (similar to Fig 2J) while in others GFP RFP cells are in an ectopic disc (Fig 3I-O, see also Fig 2 M-P). Ectopic discs were not observed at earlier time points in the same cohort of larvae, indicating that ectopic disc growth occurs between 48 and 74 h after IR, which is in agreement with our published results [14]. RFP^+^ GFP^+^ cells of the ectopic discs appear contiguous with cells of the hinge and the pleura (arrowhead and arrow, respectively, in Fig 3I-K). Of the remainder of the discs, eight resembled -IR controls and one resembled the disc shown for the 24 h time point in Fig 3A-C.

To summarize and interpret the time-course data, at 24 h after IR, GFP-marked cells of the hinge and the pleura are found in the notum but most of these cells retain RFP. At 48 h after IR, GFP^+^ cells are found deeper (more dorsal) in the notum, they are more numerous than at 24 h and most have lost RFP. At 72 h after IR, while most of the discs resemble the 48 h discs, a significant fraction showed RFP in the GFP^+^ cells in the notum. We interpret these data to mean that hinge/pleura cells translocated into the notum and lost their hinge/pleura fate, but that some of them re-gain the fate as they form ectopic discs. We cannot rule of the formal possibility that GFP^+^ RFP^+^ cells in the notum and ectopic discs at 72 h after IR formed de novo and bear no relation to the cells of the hinge and the pleura of the primary disc. But the finding that GFP^+^RFP^+^ cells in the notum are contiguous with the primary hinge and the pleura, combined with the sequence of events in the time course, make us favor the scenario of dynamic cell fate changes. In this interpretation, hinge cells that translocate into the notum originate from part the hinge that is outside the 30A domain (arrowheads in Fig 1J, L). This explains why we never saw cells of the 30A domain translocate into the notum in our previous studies [13, 14].

### Cells of the notum contribute to the ectopic disc

Confocal imaging and close inspection of each optical section showed that ectopic discs include cells that lacked both RFP and GFP (Fig 3M-O, arrows). In these experiments, only cells that lacked RFP and GFP were notum cells, suggesting that cells of the notum also contribute to ectopic discs in addition to cells that originate from the hinge. We addressed this possibility directly by lineage-tracing with a notum-specific GAL4 driver (Fig 4).

**Figure 4.**
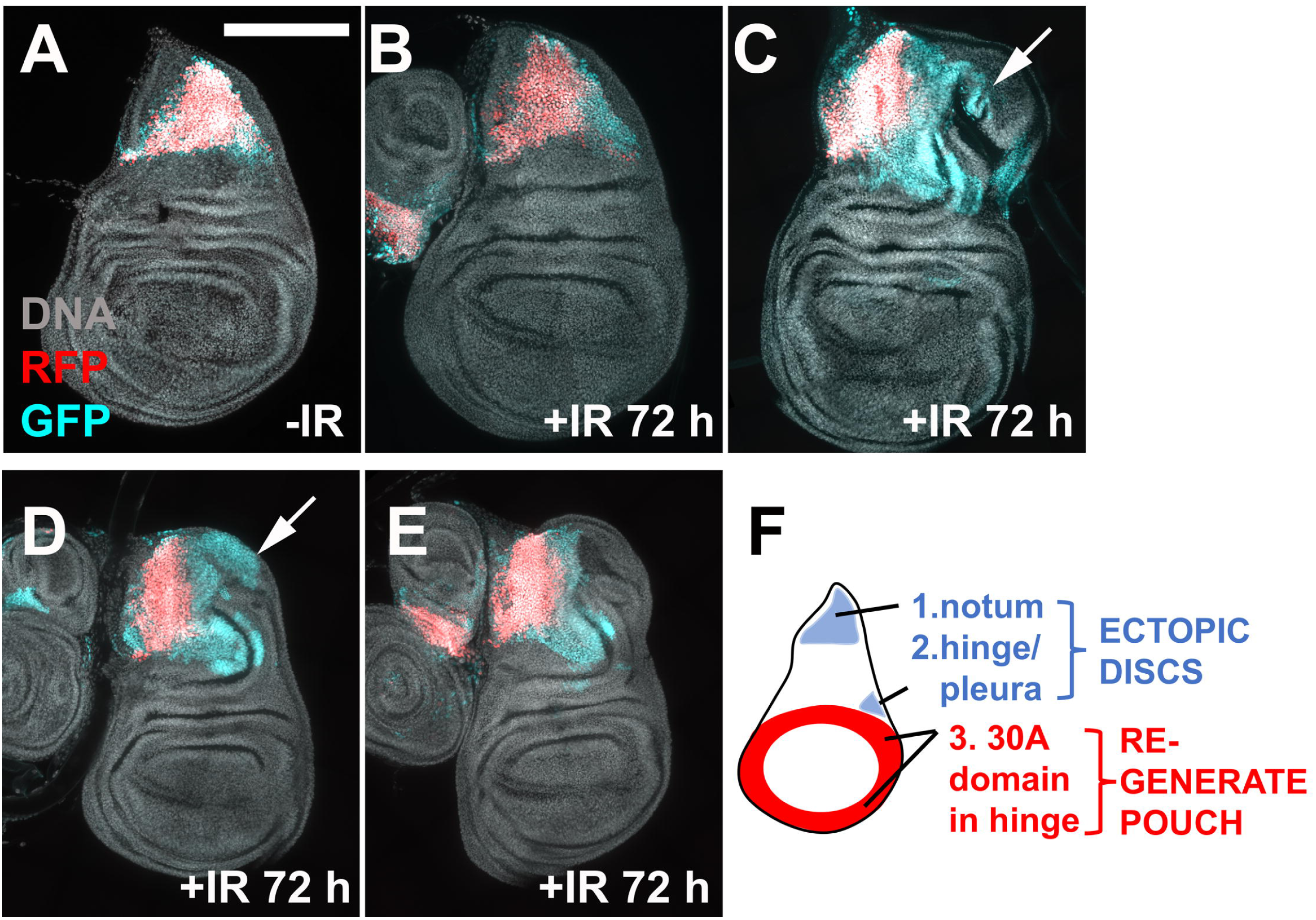
Cells of the notum contribute to ectopic discs. Larvae of the genotype UAS-G-trace/+; GAL80^ts^ / R76A01-GAL4 were treated as in Fig 1M. Wing discs are removed, fixed and imaged for RFP/GFP. The discs were also stained for DNA. All discs are shown with anterior (A) left and dorsal (D) up. Scale bar = 100 microns. Arrows = GFP^+^ RFP^-^ cells that are part of the ectopic disc.

R76A10-GAL4, bearing an enhancer fragment from the *tailup* locus, is active exclusively in a subset of notum cells (Fig 4A). Without IR, cell fate in this domain appears stable as GFP and RFP overlap (Fig 4A). At 72 h after IR, GFP/RFP overlap looks similar to –IR in most discs (Fig 4B). The exceptions are irradiated discs with ectopic growths, where we observed an expansion of the GFP cell population beyond the RFP^+^ area (Fig 4C-E). The lack of RFP in these cells suggests that they have lost their original fate as detected by R76A10-GAL4>RFP expression. Such GFP^+^ RFP^-^ cells are observed to be part of the ectopic disc, although the extent of their contributions to the ectopic disc and their location within the ectopic disc was variable from disc to discs (arrows in Fig 4C and D). Regardless, cells that originated from the notum appear to contribute to the ectopic disc in these experiments, corroborating the finding with the R73G07-GAL4 driver that ectopic discs are composed of cells that originated from the notum, the hinge (outside the 30A domain), and the pleura (modeled in Fig 4F).

In our previous studies, cells of the pouch, marked with rn-GAL4>G-trace, did not change fate or translocate after irradiation ([13]; reproduced in Fig 5A-H). Even in experiments when we directed cell death to the hinge and left the pouch cells alive, the hinge was repaired with the hinge cells and not the pouch cells [13]. We confirmed these findings using two additional pouch GAL4 drivers, R42A07-GAL4 (from the *dve* locus) and R85E08-GAL4 (from the *salm* locus). We saw little movement of pouch cells in these experiments (Fig 5I-P). Taken together, we conclude that among the cells of the wing disc, cells in the hinge, the pleura and the notum exhibit cell fate change and translocation after irradiation (Fig 4F). Of these, hinge cells in the 30A-GAL4 domain translocate and change fate in nearly all irradiated discs to regenerate the pouch [13]. In contrast, hinge cells outside the 30A domain, pleura and notum cells produce an ectopic disc in a fraction of irradiated discs.

**Figure 5.**
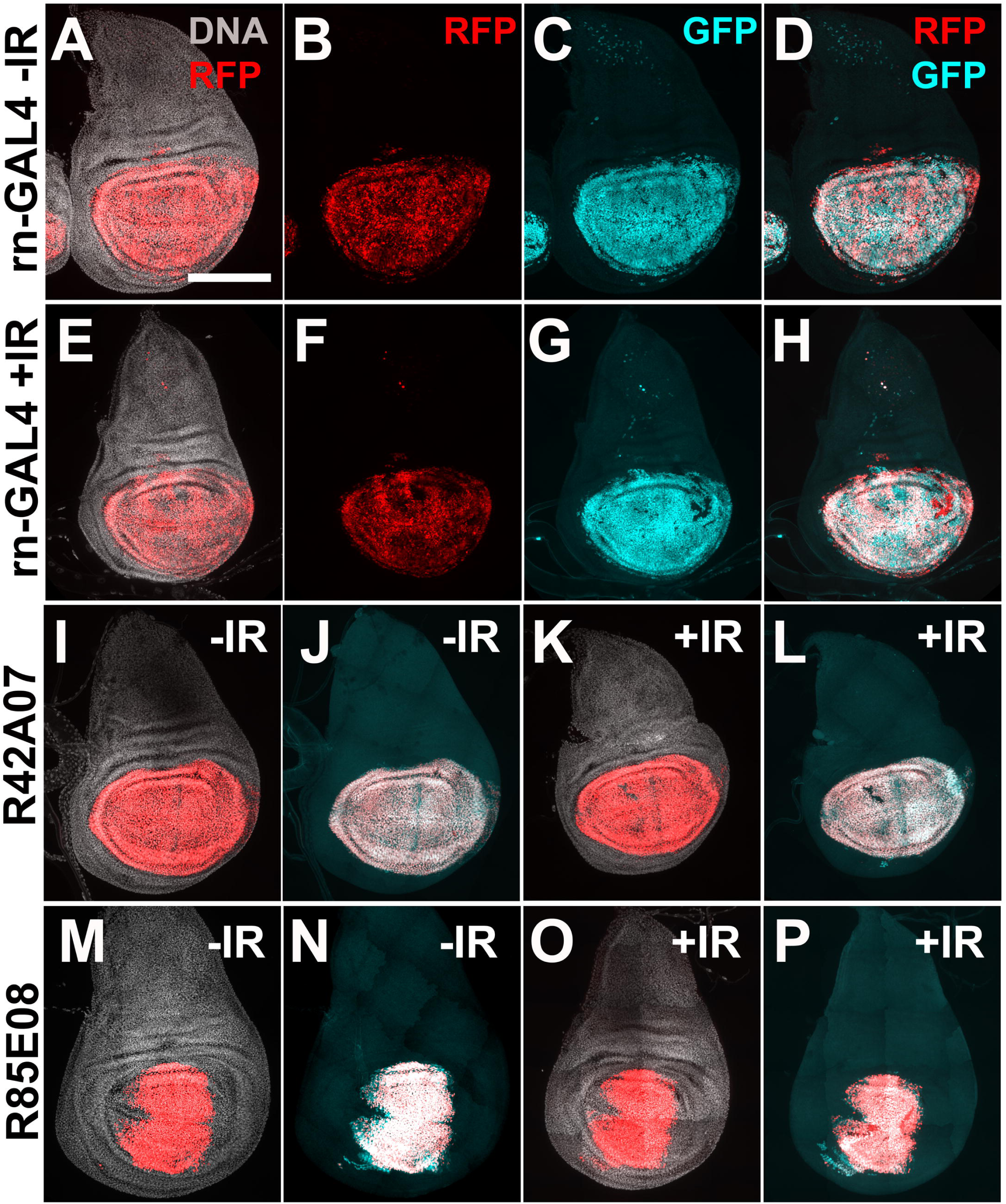
Cells of the pouch do not show regenerative properties after irradiation. Larvae of the genotype UAS-G-trace/+; GAL80^ts^ / rn-GAL4 (A-H), R42A07-GAL4 (I-L) or R85E08-GAL4 (M-P) were treated as in Fig 1M. Wing discs are removed, fixed and imaged for RFP/GFP. The discs were also stained for DNA. All discs are shown with anterior (A) left and dorsal (D) up. Scale bar = 100 microns.

### Compartment boundaries remain intact during regeneration but only in the primary disc

*Drosophila* wing disc is sub-divided into compartments, Anterior/Posterior and Dorsal/Ventral, for example, with cell lineages restricted to each compartment during development. A recent study showed that upon massive damage to the wing discs, caused by directed expression of a pro-apoptotic gene in a specific compartment, compartment boundaries collapse and are rebuilt during regeneration [24]. Further, some cells of one compartment assumed the identity of another, adjacent compartment during this process, overcoming lineage restrictions seen during development. In these models, one compartment suffered massive damage while the other compartment remained untouched. In contrast, exposure to IR causes damage that is scattered throughout the disc. To ask if compartment boundaries are breached during regeneration after IR damage, we used the same compartment-specific GAL4 drivers as in the published study, ci-, en-, and ap-GAL4, to express G-trace in the anterior, posterior and dorsal compartments of the wing disc, respectively. We used the protocol in Fig 1M to assay for breach of compartment boundaries during regeneration from X-ray damage. Without IR, ci- or en-GAL4>G-trace expressing discs show overlap of GFP/RFP (Fig 6A-D, I-J). In some ap-GAL4>G-trace discs, we observed small populations of cells that breach the boundary without IR (arrow in Fig 6R). More important, irradiated discs did not appear different from –IR control using all three compartment-specific drivers (Fig 6). This observation applies to the primary disc. In contrast, we observed fluid compartment boundaries in the ectopic disc, which may generate compartment boundaries de novo. For example, en-GAL4>RFP^+^ and RFP^-^ cells co-mingle (Fig 6M-P). Furthermore, the presence of RFP^-^ GFP^+^ cells (arrow in Fig 6P) suggests that cells that used to have the posterior identify have lost it. We conclude that pre-existing compartment boundaries remained intact during regeneration from IR damage but are more fluid in the ectopic disc.

**Figure 6.**
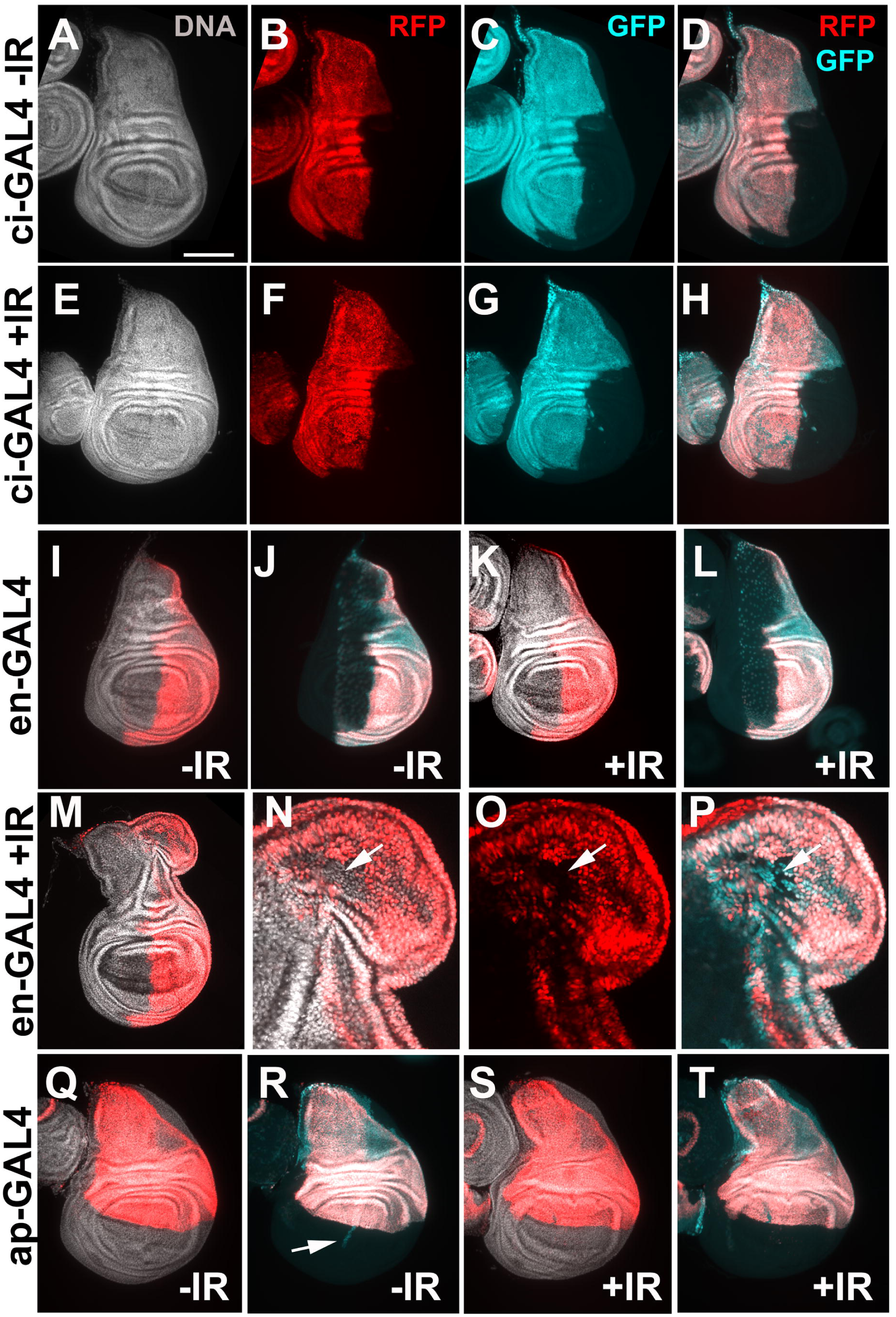
Cells do not cross pre-existing compartment boundaries during regeneration after IR. Larvae were treated as in Fig 1M. Wing discs are removed, fixed and imaged for RFP/GFP. The discs were also stained for DNA. All discs are shown with anterior (A) left and dorsal (D) up. Total number of disc examined were 56 ci-IR, 22 ci+IR, 35 en-IR, 93 en+IR, 46 ap-IR, and 88 ap+IR. Scale bar = 40 microns in N-P and 120 microns in all other panels. The genotypes were: (A-H) UAS-G-trace/ci-GAL4; tub-GAL80^ts^ /+ (I-P) UAS-G-trace/en-GAL4; tub-GAL80^ts^ /+ (Q-T) UAS-G-trace/ap-GAL4; tub-GAL80^ts^ /+

### Caspase activity and/or cell death are required for IR-induced cell fate plasticity

Regenerative cells proliferate, change fate and change location, in order to rebuild damaged tissue. We report two aspects of regenerative behavior, cell fate change and translocation.

These behaviors are not seen without IR (for example, Fig 1A and Fig 4A). Therefore, we next addressed which of the consequences of IR exposure is responsible for the induction of these aspects of regenerative behavior. IR has many effects on cells including DNA double strand breaks, cell cycle arrest by checkpoints, and apoptosis. Of these, apoptosis has been demonstrated to induce one aspect of regenerative behavior, namely proliferation of the surviving cells in a phenomenon known as Apoptosis-induced-Proliferation or AiP (reviewed in [25]). Therefore, we investigated whether apoptosis is also responsible for the induction of cell fate change and translocation after IR. We used the model of hinge cells changing fate and translocating towards the pouch because this response is seen in nearly all irradiated discs, as opposed to ectopic disc formation, which occurs in a small fraction of irradiated discs [14].

We used a chromosome deficiency that deletes three pro-apoptotic genes, H99/+, that has been shown before to delay and reduce IR-induced caspase activation and apoptosis [26]. We expressed 30A-GAL4>G-trace in this background, and saw reduced extent of fate change and translocation by the hinge cells (Fig 7, compare E to D, quantified in Fig 7I). We conclude that caspase activity and/or cell death is required for IR-induced regenerative behavior of the hinge cells.

**Figure 7.**
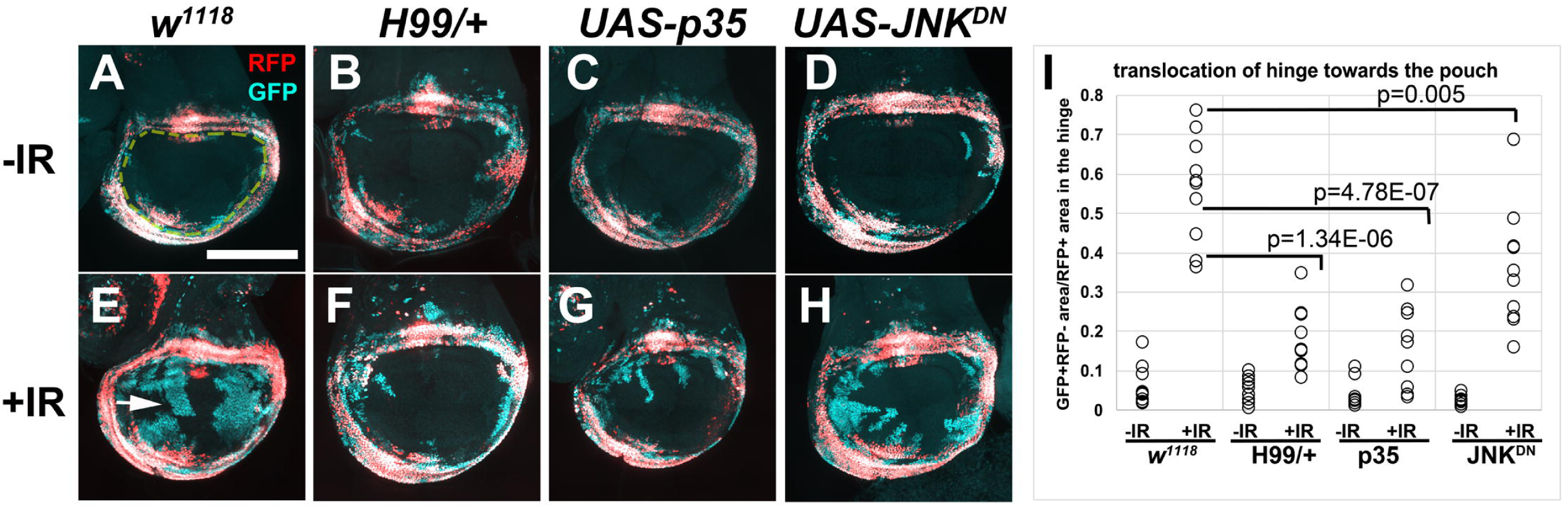
Apoptosis/caspase activity is needed for IR-induced regenerative behavior. Larvae were treated as in Fig 1M. Wing discs are removed, fixed and imaged for RFP/GFP. All discs are shown with anterior left and dorsal up. GFP^+^ RFP^-^ area within the pouch region (yellow dashed lines in A) was quantified in Image J and normalized to the RFP^*+*^ area of the hinge and plotted in (G). n=9 for H99/+ +IR and 10 each for all other samples, from two biological replicate experiments. Statistical significance was determined using 2-tailed student’s t-test. Scale bar = 100 microns. The genotypes were: W^1118^ = 30A-GAL4, UAS-G-trace; tub-GAL80^ts^ /+, from a cross of w1118 to 30A-GAL4, UAS-G-trace/CyO-GFP; tub-GAL80^ts^ / tub-GAL80^ts^ H99/+ = 30A-GAL4, UAS-G-trace/+; tub-GAL80^ts^ /H99 deficiency UAS-p35 = 30A-GAL4, UAS-G-trace/+; tub-GAL80^ts^ /UAS-p35 UAS-JNK^DN^ = UAS-JNK^DN^ /+; 30A-GAL4, UAS-G-trace/+; tub-GAL80^ts^ /+

### Effector caspase activity is required in the regenerative cells

In many models of regeneration in larval wing discs, dying cells are the source of signals that promote regenerative behavior in surviving cells. Some of these signals are produced in response to apical caspase activity in the dying cells [25]. But no study we are aware of has addressed the need for caspase activity in the regenerative cells. Yet, there is mounting evidence for the role of caspases in cell fate changes during development [27, 28]. Our identification of regenerative cells in the hinge and the ability to target UAS-transgenes to these using 30A-GAL4 allow us to directly address this possibility. To this end, we co-expressed p35, an inhibitor of effector caspases, with 30A-GAL4. We found that this treatment inhibited the translocation and fate change of the hinge cells. We conclude that caspase activity/apoptosis is needed in the hinge for regenerative behavior. We note that the hinge cells are protected from IR-induced caspase activation and apoptosis by the action of Wg and STAT [13]. The effect of a caspase inhibitor p35 on these cells suggest that caspases may be playing a non-apoptotic role in these cells (further discussed in DISCUSSION). JNK kinases are stress-responsive; in *Drosophila* wing discs, JNK acts in the dying cells to produce mitogenic signals, which then stimulate neighboring surviving cells to proliferate [25]. To address the role of JNK in *regenerative* cells, we co-expressed a previously characterized dominant negative JNK using a UAS transgene [29]. We find that JNK^DN^ reduced the translocation and fate change of the hinge cells, but its effect was not as severe as inhibition effector caspases (Fig 7D-H, quantified in Fig 7I).

## DISCUSSION

In tumor biology, the concept of cancer stem cells has been controversial, but there is agreement that within a tumor, some cancer cells are better than others at re-initiating tumor growth [15, 16]. Such ‘Cancer Stem-like Cells’ (CSCs) are recognized by specific protein markers and by their greater ability, relative to cells without such markers, to form tumor spheres in culture or new tumors in mice. Eradication of CSCs is considered necessary for successful therapy. However, not only do CSCs generate non-stem cancer cells, non-stem cancer cells also are capable of converting to CSCs. Even more concerning, cancer treatments including IR converts non-stem cancer cells from a variety of cancer types into cells with CSC markers that can initiate new tumors in culture and in vivo [19–21]. An estimated 50% of cancer patients receive IR, alone or as part of their treatment (www.cancer.org). Yet, we know very little about what aspects of IR exposure induce CSCs. The finding that IR induces non-stem cells of the *Drosophila* larval wing discs to exhibit stem cell like properties allows us to fill in the gaps in this knowledge.

One key gap concerns the question ‘what are the consequences of IR that induce stem cell-like behavior?’ The answer, we report here, is caspase activity. Surprisingly, we detect this requirement in regenerative cells that translocate and change fate. Regeneration typically relies on surviving cells proliferating and re-programming to replace cells lost to cell death or surgical removal. Prior work on *Drosophila* wing discs found that cell death itself induces proliferation of the surviving cells (AiP). In this response, signaling through apical caspase Dronc and JNK in dying cells act through Dpp and Wg to promote cell division in the surviving cells [25]. This and other similar mechanisms that operate in other larval discs explain the proliferative aspect of regeneration, but the re-programming aspect remained to be better understood. Our study fills the gap in the knowledge by identifying the role of effector caspases. We note two key differences between AiP and fate change/translocation. The former requires apical caspases but not effector caspases [25], while the latter requires effector caspases (this report). Further, the requirement for effector caspases is within the regenerative cells.

There are several insightful reports of regeneration after genetic ablation where cell death is directed to a specific wing compartment (for example, [30–33]). Comparison of results from these studies with ours identify similarities as well as differences. Regeneration of the pouch by the hinge after IR damage we study resembles the model in which rn-GAL4 drove the expression of pro-apoptotic Hid to kill the pouch cells [30]. Lineage tracing in this model showed that the pouch was repaired, at least in part, by relocating hinge cells. The loss of hinge fate was not monitored in this study. Compartment boundaries provide another example of similarities/differences between different models. Compartment boundaries were breached during regeneration after genetic ablation of some wing disc compartments (A/P; [24]) but not others (pouch; [32]). In the latter case, mutations in Taranis, which is required to reestablish engrailed expression in the regenerated posterior cells, did cause the breach of compartment boundaries. We find intact compartment boundaries in regenerated primary disc after IR, but fluid boundaries in the ectopic disc. This contrasts with a report that ectopic discs that form after genetic ablation of the pouch shows intact compartment boundaries that are continuous with the pre-existing ones [33]. We note that even in genetic ablation models, the choice of apoptotic gene used to kill cells can have different outcome on the regenerative behavior of the surviving cells. For example, ablation of the pouch using Eiger produced ectopic discs but ablation of the pouch using Rpr did not [33]. Collectively, these data illustrate how different regenerative models rely on different cellular behaviors, hence the need to study each to learn the range of possibilities. This, we believe, is particularly true in the case of IR where damage (i.e. cell death) is not confined to any particular compartment but scattered throughout the wing disc in a reproducible pattern [13].

We report the identification of two additional pools of cells that exhibit IR-induced fate change and translocation, bringing the total to three (Fig 4F). A recent study used genetic ablation to probe the regenerative potential of the notum. After ablation of the pouch and the hinge, the notum showed no increase in proliferation, did not regenerate the hinge or the pouch, and instead duplicated itself. The authors concluded that the notum cells have little regenerative potential [31]. This is in sharp contrast to IR-induced regenerative behavior we see in the notum. After IR, we detect a 3-fold increase in mitotic activity in the notum [14], and lineage tracing shows the notum contributes to the ectopic discs (this report). Our results parallel more closely what happens after genetic ablation of the pouch in a recent study, which resulted in the production of ectopic wing discs in some mutant backgrounds [33]. Lineage tracing suggested that notum cells changed fate to contribute to the ectopic disc, which is perfectly in agreement with our results. We add to this picture by identifying additional pools of cells in the dorsal-posterior hinge/pleura that contribute to the ectopic discs.

We identified a requirement for effector caspase activity in the hinge cells that change fate and translocate towards the pouch. Apical caspases such as Dronc in *Drosophila*, are known to provide non-apoptotic functions, for example in initiating pro-proliferative signals to neighboring survivors in AiP [34–37]. These functions occur in addition to Dronc’s role in activating effector caspases and apoptosis. What we describe here suggests a non-apoptotic role of effector caspases, in cell fate changes. There is precedent for non-apoptotic roles of *effector* caspases ([36, 38–40]; reviewed in [27]). In a particularly relevant study, effector caspase CED-3 in *C. elegans* was found to cleave three cell fate determinants encoded by *lin-28*, *disl-2*, and/or *lin-14*, all of which are unrelated to apoptosis, in order to ensure that cell fate changes and developmental transitions occur normally [28]. IR induces caspase activation and apoptosis by inducing IAP antagonists Hid and Rpr. After irradiation, *hid* and *rpr* are transcriptionally induced throughout the wing disc except in the hinge [13]. In the hinge, we found previously that *hid* transcription is induced but *rpr* is repressed in a Wg-dependent manner [13]. We further demonstrated that *rpr* repression can explain the observed resistance to IR-induced apoptosis in the hinge. Based on the current findings, we suggest that IR activates caspases in hinge cells but insufficiently for apoptosis. Instead, non-lethal levels of effector caspase activity, we propose, result for IR and act to facilitate two aspects of regenerative behavior in these cells, fate change and translocation towards the pouch. If true, partial effect of JNK^DN^ we see be explained by a recently documented role of JNK [41]. This study found that in a *Drosophila* model of oncogenic RAS-driven tumors, JNK participates in a positive feed-back loop to amplify caspase activity. Caspases in this context is non-apoptotic but instead promote tumor growth and invasion. We speculate that JNK may also help amplify or sustain non-apoptotic caspase activity in the hinge cells of the 30A domain, which can explain why regenerative inhibition of JNK curbs their regenerative behavior.

Finally, cells of the pouch displays little indication that they change fate or translocate (Fig 5) and not even when we directed cell death to the hinge and left the pouch cells alive [13]. This parallels how IR induces Cancer Stem Cell-like properties in some cancer cells but not others, a phenomenon for which we lack a mechanistic understanding. Identification of distinct pools of cells with different abilities to respond to IR in *Drosophila* will allow the identification of underlying mechanisms in the future.

## MATERIALS AND METHODS

### Drosophila stocks and methods

These stocks are described in Flybase: *w^1118^*, 30A-GAL4 (on Ch II, Bloomington stock# or BL37534), Ptub-GAL80^ts^ (on Ch III), *rn-GAL4* (on Ch III), *en-GAL4* (on Ch II), *ci-GAL4* (on Ch III), *ap-GAL4* (on Ch II), UAS-p35 (on Ch III), and UAS-JNK^DN^ (on Ch I, BL6409). The stock used for lineage tracing is also described in Flybase; w^*^; P{UAS-RedStinger}4, P{UAS-FLP.D}JD1, P{Ubi-p63E(FRT.STOP)Stinger}9F6 /CyO (BL28280). Genotypes for some BL stocks are in S1 Table and include FlyLight stocks [16]. For all experiments except Fig 7, G-trace/CyO-GFP; GAL80^ts^/GAL80^ts^ virgin females were crossed to males with various GAL4 drivers. For Fig 7, 30A-GAL4>UAS-RFP, G-trace/CyO-GFP; GAL80^ts^ /GAL80^ts^ virgin females were crossed to *w^1118^* males (*w^1118^* controls) or H99/TM6-TB males or UAS-p35/UAS-p35 males. For JNK^DN^ experiments, UAS-JNK^DN^ homozygous virgin females were crossed to 30A-GAL4>UAS-RFP, G-trace/CyO-GFP males. Progeny bearing G-trace (RFP^+^ GFP^+^ larvae) were sorted for use.

### Larvae culture and irradiation

Larvae were raised on Nutri-Fly Bloomington Formula food (Genesee Scientific) according to the protocol shown in Fig 1M. The cultures were monitored daily for signs of crowding, typically seen as ‘dimples’ in the food surface as larvae try to increase the surface area for access to air. Cultures were split at the first sign of crowding. Larvae in food were placed in petri dishes and irradiated in a Faxitron Cabinet X-ray System Model RX-650 (Lincolnshire, IL) at 115 kv and 5.33 rad/sec.

### Antibody staining

Antibodies to Zfh2 (1:400, rat polyclonal, [42]) and anti-rat secondary antibodies (1:200, Jackson) were used. In all experiments, wing discs were dissected in PBS, fixed in 4% para-formaldehyde in PBS for 30 min, and washed three times PBTx (0.1% Triton X-100). For antibody staining, the discs were washed in PBS instead of PBTx after the fixing step, permeabilized in PBTx with 0.5% Triton X-100 for 10 min and rinsed in PBTx. The discs were blocked in 5% Normal Goal Serum in PBTx for at least 30 min and incubated overnight at 4°C in primary antibody in block. The discs were rinsed thrice in PBTx and incubated in secondary antibody in block for 2 h at room temperature. Stained discs were washed in PBT. The discs were counter-stained with 10 μg/ml Hoechst33258 in PBTx for 2 min, washed 3 times, and mounted on glass slides in Fluoromount G (SouthernBiotech).

### Image Analysis

With the exceptions noted below, the discs were imaged on a Perkin Elmers spinning disc confocal attached to a Nikon inverted microscope, using a SDC Andor iXon Ultra (DU-897) EM CCD camera. The NIS-Elements acquisition software’s large image stitching tool was used for the image capture. 15 z-sections 1 um apart were collected per disc. Sections that exclude the peripodial cells were collapsed using ‘maximum projection’ in Image J. The exceptions are images in S1 Fig which were acquired on a Leica DMR compound microscope using a Q-Imaging R6 CCD camera and Ocular software.

### Statistical Analysis

For sample size justifications, we used a simplified resource equation from [43]; E = Total number of animals − Total number of groups, where E value of 10-20 is considered adequate. When we compare two groups (*w^1118^* vs H99/+, for example), 6 per group or E = 11 would be adequate. All samples subjected to statistical analysis exceed this criterion. Two tailed student t-tests were used in Fig 7.

## ACKNOWLEDGMENTS

*Drosophila* stocks from the Bloomington *Drosophila* Stock Center (NIH P40OD018537) were used in this study. We are grateful to the FlyLight team for generating and distributing these stocks, without which this study would not be possible. We thank Chris Doe for anti-Zfh2 antibodies. We thank the Light Confocal Microscopy Facility of MCD Biology, CU Boulder, for assistance with imaging.

---

**S1 Figure. Lineage tracing with Flylight GAL4 drivers**

Larvae were treated as in Fig 1M –IR. Wing discs are removed, fixed and imaged for RFP/GFP. All discs are shown with anterior (A) left and dorsal (D) up. Scale bar = 120 microns. The genotypes were: UAS-G-trace/+; tub-GAL80^ts^ /GAL4. Bloomington stock number (BL) and Flylight construct number (R-) and the locus of origin for the enhancer are indicated on each panel. We also tested and found no RFP/GFP expression with *BH-1* R81E08 (BL40117) and doc-1 R45H05 (BL46529), and weak RFP expression in the peripodium with <u>unc5</u> R93E10 (BL48420). R85E08 (*salm*), R42A07 (*dve*) and R76A01 (*tup*) showed good overlap of RFP/GFP and were used in lineage tracing studies.

**S2 Figure. G-trace expression in cells outside the columnar epithelial layer**

Wing discs were removed from 3^rd^ instar larvae expressing R73G07-GAL4>G-trace and treated as in Fig 1M-IR, fixed and imaged for GFP. The disc shown is the same as in Fig 2A-D. Three optical sections illustrate peripodial cells (A, arrows), columnar epithelial cells (B, but also visible in other optical sections) and tracheal cells (C, arrows). The disc is shown with anterior (A) left and dorsal (D) up.

**S1 Table. Genotypes and stock numbers for stocks from Bloomington Stock Center used in this work.**

